# A Parallel Accumulation-Mobility Aligned Fragmentation Strategy Utilizing High-Resolution Ion Mobility for High Performance Proteomics Analysis

**DOI:** 10.64898/2026.02.09.704896

**Authors:** Leonard C. Rorrer, Liulin Deng, Lauren Royer, Isabel Uribe, Benjamin C. Orsburn, Oliver Bernhardt, Tejas Gandhi, Lukas Reiter, Daniel DeBord

**Affiliations:** MOBILion Systems, Inc. 4 Hillman Drive, Suite 130, Chadds Ford, PA 19317, United States; Organ Pathobiology and Therapeutics Institute, University of Pittsburgh, 3501 5th Avenue, Pittsburgh, PA 15260, United States; Biognosys, Wagistrasse 21, 8952 Schlieren, Switzerland

## Abstract

Here we present a novel data independent acquisition (DIA) mass spectrometry (MS) operating mode termed parallel accumulation-mobility aligned fragmentation (PAMAF) that offers enhanced speed and sensitivity of ion fragmentation analysis for nontargeted discovery workflows such as bottom-up proteomics. This mode of operation leverages high-resolution ion mobility (HRIM) separation capabilities of the structures for lossless ion manipulation (SLIM) technology to achieve HRIM-based precursor isolation in place of traditional quadrupole filtering approaches. This PAMAF mode of operation increases the number of features that can be identified per MS1/MS2 acquisition cycle by employing mobility-based time alignment to associate fragment ions with their corresponding precursor ions. By using a high-speed, lossless separation technique for precursor isolation instead of the comparatively slow and wasteful quadrupole filtering method, we can avoid ion losses up to 99% while simultaneously increasing the rate at which precursor ions are sequentially fragmented and detected. Additionally, by storing ions in a trapping region while the previous packet of ions is being analyzed, the PAMAF mode achieves ∼100% ion utilization efficiency. Benchmarking results of LC-PAMAF-MS analysis of a whole cell protein digest showed approximately 6x more protein group identifications compared to a standard data-dependent acquisition (DDA) analysis without HRIM on the same QTOF instrument, and to over 100x improvement for low-load workflows. Quantitative evaluations demonstrated that PAMAF mode could quantify low abundance peptides, including those undetectable by DDA. Additionally, since precursor isolation in PAMAF mode is size-based rather than *m/z*-based, many coeluting isobars and isomers can be resolved prior to fragmentation to eliminate chimeric spectra that compromise identification accuracy. In this work we also explored the benefits of combining HRIM and quadrupole isolation to achieve improved specificity. This approach, known as DIA-PAMAF mode, further reduces the frequency of chimeric fragmentation spectra, and enabled the detection of over 8,000 protein groups from a HeLa digest analysis. PAMAF mode brings a powerful new technique to the field of proteomics that has the potential to improve the sensitivity and selectivity of mass spectrometry-based proteomics.

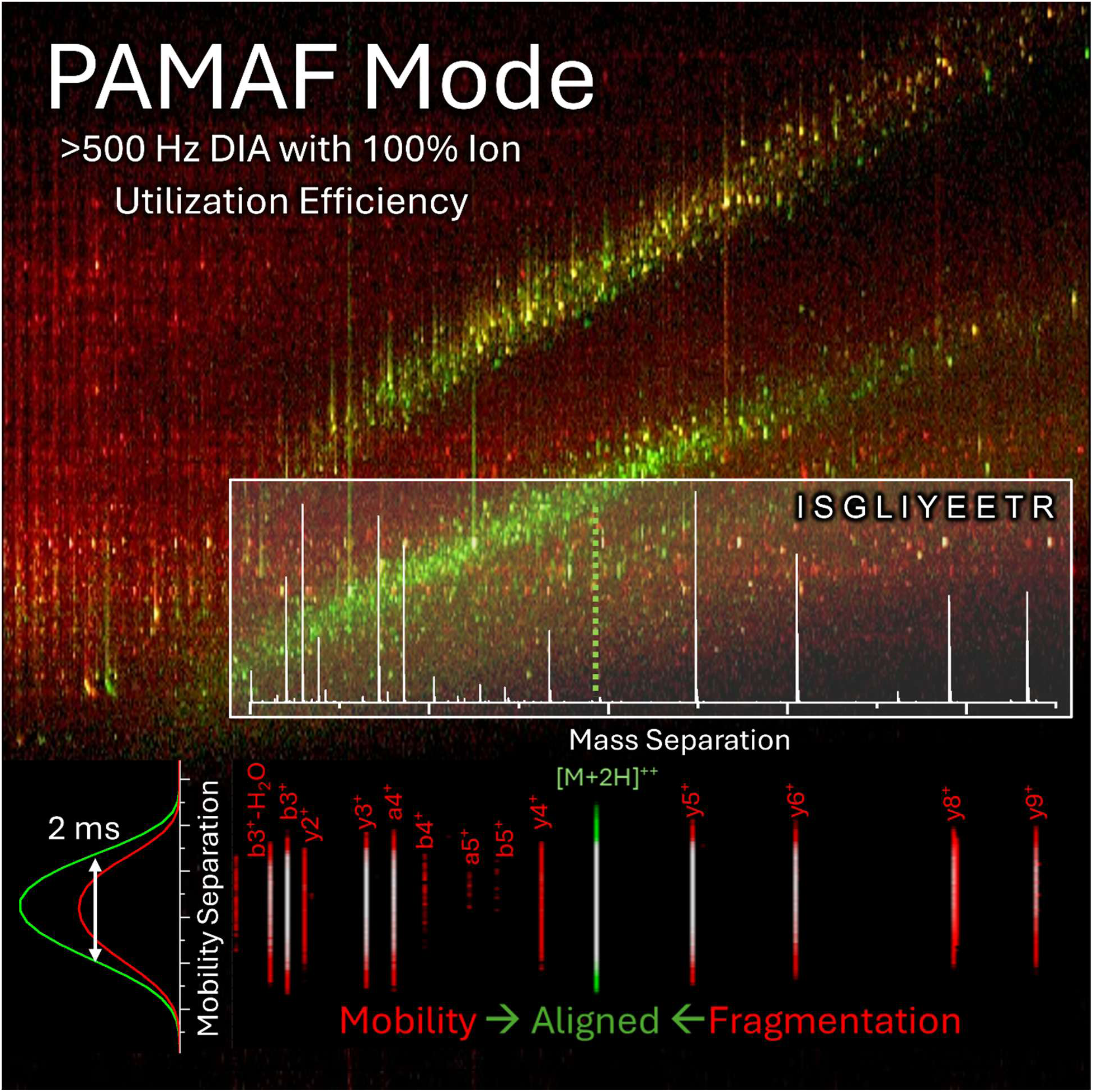

## INTRODUCTION

High-performance mass spectrometry has become the gold standard for molecular analysis of complex biological systems, particularly in proteomics research^1–3^. The most commonly employed modes of operation rely on tandem mass spectrometry, or “MS/MS” to achieve the necessary specificity of detection within these complex matrices^1,4,5^. At the simplest level, tandem mass spectrometry is the process by which an ion (precursor ion) is fragmented into pieces (fragment ions) that are representative of the precursor ion. In bottom-up proteomics, precursor ions are typically peptides from a digested protein or protein mixture and the fragment ions from these peptides are a sort of barcode which can be used to positively identify the original peptide.

While the use of MS/MS for proteomic analysis provides a very powerful, highly specific tool which has seen steady growth of implementation, there can be drawbacks associated with this technique. To generate the highest quality and most representative fragmentation spectrum for a given peptide ion, ions other than the precursor of interest are typically discarded by means of a quadrupole mass filter which transmits a selected ion or range of ions while discarding other ions outside of the desired range. This process can be done in either a data-dependent manner (data-dependent analysis, DDA), where only specific precursor ions of interest that meet threshold criteria are transmitted and fragmented, or in a data-independent manner (data independent analysis, DIA), where quadrupole isolation steps through pre-determined ranges of precursor ion isolation windows^6,7^. Both have their benefits, with DDA generally offering high quality fragmentation spectrum for a limited number of selected peptides, and DIA generally offering lower quality fragmentation spectrum for a greater number of peptides in a more reproducible way. Both DDA and DIA often perform multiple sequential MS2 “isolation events” between MS1 scans to provide coverage of the full range of desired precursor ions. Dividing the cycle time among multiple MS2 scans, together with quadrupole filtering that discards all but a small subset of ions, reduces the system’s effective ion utilization efficiency and can significantly impact sensitivity. Quadruple mass filtering prior to fragmentation is also blind to the presence of isobaric or isomeric species which would be transmitted/fragmented simultaneously within a given mass isolation window, thereby generating chimeric fragmentation spectra and leading to convolution of the identification for that specific peptide. This is often most apparent in cases of peptides that may be modified at two or more potential sites^8^.

The wasteful nature of quadrupole-based MS/MS was first recognized and a solution proposed nearly 25 years ago^9,10^. This method introduced by the Clemmer lab was called parallelized collision-induced dissociation (Parallel CID)^11^. In this method, drift tube ion mobility separation was used to separate ions prior to fragmentation, allowing fragment ions to appear at parallel drift times with their corresponding precursors, thereby enabling straightforward association and direct generation of fragmentation spectra for all precursor ions without requiring a quadrupole filtering step^12–14^. This approach proved successful for the analysis of simple peptide mixtures, but as noted by the authors, application to more complex samples proved impractical given the limited resolving power (20-30) achievable of their prototype IM-CID-TOF system. For reference, typical DIA workflows employ quadrupole isolation windows of 2-20 Th wide for the analysis of tryptic peptide ions span a *m/z* range of 600 Th (from ∼400 Th to 1,000 Th). This means the population of peptide ions are effectively filtered with a resolving power of 30-300, setting the bar for the level of specificity required if ion mobility is to be employed to replace the functionality of quadrupole precursor isolation^15,16^.

The concept of ion mobility-enabled ion fragmentation analysis was first commercialized by Waters in the form of their High Definition Mass Spectrometry (HDMS^E^)^17^ and Ultra Definition Mass Spectrometry (UDMS^E^)^18^ operating modes employed on the Synapt series of traveling wave ion mobility spectrometry (TWIMS) QTOF mass spectrometers. Application to proteomics analyses was explored, detecting up to ∼5400 protein groups from a human cell lysate in a 180 minute LC analysis, but again, the limited peak capacity of the IM technology capped achievable performance^18^. Ultimately, it was not until the introduction of Parallel Accumulation Serial Fragmentation (PASEF) mode by Bruker^19^ on their line of timsTOF mass spectrometry systems that ion mobility-enabled proteomics was demonstrated to be superior to conventional MS/MS-based approaches of the day. However, the various PASEF modes of operation still rely on combining quadrupole filtering with trapped ion mobility spectrometry (TIMS) to overcome the peak capacity limitation still present with TIMS. Only through this combination of TIMS-quadrupole filtering is the necessary level of isolation specificity for high performance proteomics achieved on these systems, but this approach retains some of the original limitations of quadrupole filtering outlined above.

This current work expands upon the original concept of “Parallel CID” by employing a next-generation HRIM technology leveraging SLIM^20^. Introduced in 2014, SLIM is the only ion mobility technology developed to date that is capable of providing an ion mobility resolving power on par with a quadrupole-based DIA workflow across the full range of peptides in a single scan^20–22^. It does so by enabling significantly longer separation path lengths (e.g. 13 m), owing to its printed circuit board-based electrode design that allows the ion path to be folded back on itself in a serpentine arrangement. This results in collision cross section (CCS) based ion mobility resolving powers (R_P_) of 250-300 (CCS/ΔCCS)^23^, which means the SLIM based HRIM separation is capable of direct precursor isolation within a proteomic mixture without requiring additional quadrupole filtering.

This approach provides a means to quickly and efficiently isolate precursor ions prior to fragmentation and detection. HRIM isolates ions in time using a high-speed (i.e. <1 sec), gas phase separation contrary to a quadrupole filter which discards non-selected ions. This HRIM separation occurs prior to the fragmentation step in MS/MS analysis, which allows for mobility alignment of precursor ions observed in the high-resolution mobility spectrum to be mapped back to the fragment ions observed in the subsequent mobility spectrum as they are aligned in the mobility space. This mobility-aligned fragmentation, or MAF^24^, produces alternating ion-mobility-mass spectrometry data frames corresponding to a frame with no collision energy (MS1) and a data frame with collision energy (MS2). The SLIM technology also allows for much larger ion populations to be accumulated and stored prior to analysis, with capacities of over 10^8^ stored charges being demonstrated, which is 10-100x higher than other ion trapping technologies^25^. Greater charge capacity results in the ability to increase sensitivity and dynamic range of analysis. The sensitivity of analysis is further expanded to maximize ion utilization through parallel accumulation (PA), where ions can be accumulated while simultaneously separating a previous collection of ions. In conjunction with the MAF described above, a new parallel accumulation mobility aligned fragmentation (PAMAF) mode has been developed, tested, and is described herein.

## EXPERIMENTAL PROCEDURES

### Samples

For benchmarking the performance of the PAMAF operation mode, a variety of standardized samples were used. These samples included the Pierce Peptide Retention Time Calibration mixture (PRTC, P/N 88320, Thermo Fisher, Waltham, MA), Pierce HeLa Protein Digest Standard (P/N 88328, Thermo Fisher, Waltham, MA), MS Qual/Quant QC Mix (P/N MSQC1, Millipore Sigma, St. Louis, MO), and SpikeMix™ Kinase Activation Loops–phosphorylated mixture (P/N SPT-KAL-POOL-L-10pm-phospho, JPT peptide Technologies, Berlin, Germany). The PRTC sample contains 15 stable isotope labeled peptides and was received as a 0.5 µM solution. The solution was either used as received or diluted further via serial dilution in water with 0.1% formic acid to a final desired test concentration. The HeLa sample contains a mixture of trypsin-digested proteins and was received as a lypholized powder and reconstituted in water with 0.1% formic acid and serially diluted to a final concentration dependent on sample load and experiment needs. For samples where the HeLa concentration was less than 10 ng/µL, 0.015% w/v n-Dodecyl β-D-maltoside (DDM) was added as a surfactant to prevent loss of material to sample vial walls. Additionally, to aid in retention time calibration for the HeLa samples analyzed by liquid chromatography, Indexed Retention Time calibration peptides were added (Biognosys iRT Kit, P/N 1816351, Bruker, Billerica, MA). The MS Qual/Quant sample contains a mixture of six trypsin*-*digested proteins, and 14 stable-isotope labeled peptides, corresponding to 2-3 peptides from each of the digested proteins. The sample was received as a lypholized powder, dissolved into water with 0.1% formic acid, and serially diluted to final test concentrations with the same water/0.1% formic acid solvent. The phosphorylated peptide standard contains 288 phosphorylated, stable isotope labeled peptides from human kinase activation loops. The sample was received as a lypholyzed powder, dissolved into water with 0.1% formic acid, and serially diluted to final test concentrations.

### Liquid Chromatography

For samples analyzed by liquid chromatography, an Evosep One UHPLC (Evosep, Odense, Denmark) was used. Samples were prepared on the Evotips following the vendor instructions exactly. All analyses with Evosep used the standard, preconfigured Evosep methods with no modifications. Mobile phases A and B were water with 0.1% formic acid (v/v) and 100% acetonitrile with 0.1% formic acid (v/v), respectively. For 30 samples per day (SPD) methods, samples were separated on a 15 cm x 150 µm i.d. reversed-phase column packed with 1.5 µm C_18_-coated porous silica beads heated to 40°C (Evosep EV 1137)^26^. For 200 SPD methods, samples were separated on a 4 cm x 150 µm i.d. reversed-phase column packed with 1.5 µm C_18_-coated porous silica beads (EV1109)^27^. For low-load studies, the Evosep system was configured in the Zoom mode to run the Whisper Zoom methods. For the 40 SPD Whisper Zoom methods, samples were separated on a 15 cm x 75 µm i.d. reversed-phase column packed with 1.9 µm C_18_-coated porous silica beads heated to 40°C (EV1112)^28^. For 80 and 120 SPD Whisper Zoom methods, samples were separated on a 5 cm x 75 µm i.d. reversed-phase column packed with 1.7 µm C_18_-coated porous silica beads heated to 50°C (IonOpticks Aurora Rapid, Collingwood, Australia)^29,30^. The outlet of the LC columns (with the exception of the Ion Optiks column) was connected to an Evosep EV1116 Agilent Sleeve Adapter fitted with a 30 µm stainless steel nanospray emitter (Evosep EV1086) mounted in a custom holder and positioned in front of the capillary inlet of the high-resolution ion mobility spectrometer/quadrupole time-of-flight mass spectrometer (HRIM/Q-TOF MS) described below. For analyses using the IonOpticks column, the integrated nanospray emitter was positioned directly in front of the capillary inlet using an in-house custom holder and heater.

### High-Resolution Ion Mobility/Quadrupole Time-of-Flight Mass Spectrometer

In the experiments described here, a prototype of HRIM/Q-TOF MS was used for analysis. The HRIM spectrometer is based on a MOBIE^®^ system (MOBILion Systems, Chadds Ford, PA) and was coupled to one of two Agilent Q-TOF mass spectrometers (6546 or 6545, Agilent, Santa Clara, CA). Ions produced by nanoelectrospray ionization in the inlet region of the mass spectrometer entered the vacuum region of the instrument through a glass capillary and passed through a dual-narrow ion funnel (DNIF) to collect and focus ions into the inlet of the HRIM module. Ions that enter the ion mobility region are confined between printed-circuit boards (PCBs) by RF electrical fields. A phase-shifted AC waveform (i.e. traveling wave) applied to a repeating sequence of PCB electrodes moves ions to an on-board accumulation (OBA) region where they are accumulated for some period prior to ion mobility separation. Ions are released into the long serpentine separation path while the next set of ions is accumulated in parallel. Ions are moved through the separation path with a traveling wave voltage that varies in both frequency and amplitude during the mobility separation to improve resolving power across the mobility range^31^. After mobility separation, ions are guided into the Q-TOF for either precursor mass analysis or collision-induced dissociation (CID) followed by fragment mass analysis, producing alternating ion mobility frames of low collision energy (precursor ion detection) and ion mobility frames with high collision energy, detecting fragment ions corresponding to the precursor ions from the previous frame. Optionally, ions can be filtered according to their *m/*z by using the quadrupole mass filter for pseudo-DIA-PAMAF mode experiments (describe in more detail below). For experiments where no ion mobility separation is desired, the ion mobility portion of the instrument can be configured to allow all ions to pass through without accumulation, enabling standard Q-TOF mass analysis without mobility separation. Passthrough operation was used to collect data-dependent acquisition (DDA) data used as an instrumental baseline. After mass analysis, detected ion current was digitized and converted into mass spectral data, which were saved as result files with relevant information including the LC retention time, HRIM arrival time, mass-to-charge (*m/z)*, and intensity for all data points.

### Data Processing

Data processing to identify and quantify the protein groups and peptides found within the analyzed samples was performed as follows. For DDA data, the data were first converted into .mzml format with MSConvert^32^ before processing with FragPipe v22.0^33^. The HeLa FASTA file was downloaded from NCBI on November 18, 2024 and iRT peptide sequences, decoys, and contaminants were manually added. Searches were performed with trypsin digestion (maximum allowed missed cleavages set to 2), precursor mass tolerance set to 20 ppm, fragment mass tolerance set to 25 ppm, methionine oxidation (15.9949 amu) and protein N-terminal acetylation (42.0106 amu) set as variable modifications, and cysteine carbamidomethylation (57.02146 amu) set as a fixed modification. Label-free quantification was performed with IonQuant and match between runs (MBR) was enabled with default settings. For PAMAF mode data, the data were processed with default settings using Spectronaut v19 (Biognosys, Schlieren, Switzerland)^34^. For both software packages, the same database FASTA file was used, and processing parameters were matched as close as possible (i.e. missed cleavages, minimum amino acids per peptide, number of post-translation modifications (PTMs), false-discovery rate (FDR), etc.). For some datasets, data were processed using a custom library developed using a fractionated sample dataset (Supplemental Figure S1). Data were also processed and reviewed using Skyline Targeted Mass Spectrometry Enviroment^35^ in some cases.

### Experimental Design and Statistical Rationale

Samples were grouped by mass spectrometric and liquid chromatography methods. A minimum of 3 replicate injections were performed for liquid chromatographic methods to assess the technical reproducibility of the respective methods. Dilution series were measured from low to high concentration with blanks analyzed prior to avoid carryover from previous analyses. For all data processing using FDR based identification, the FDR was set to 0.01 (1%). This study does not draw any biological conclusions.

## RESULTS

### PAMAF Mode

To evaluate the potential of the PAMAF operation mode, on a SLIM-QTOF instrument to perform “quad-free” DIA proteomics analysis, a modified MOBIE HRIM system with a 13 meter ion mobility path coupled to an Agilent 6546 or 6545 QTOF was used. **Figure 1** gives a schematic overview of the system. Samples for performance evaluations were introduced to the system using UHPLC (**Fig. 1A**) generating a chromatographic separation of the sample. After ionization, samples entered the HRIM module (**Fig. 1B**) where ions were initially accumulated in an ion trapping region prior to ion mobility separation. The system can be operated in a parallel accumulation (PA) mode where ions can be accumulated while the previously accumulated packet of ions is being separated. The user-adjustable ion accumulation time is preferably configured so that the ion accumulation and separation times are equivalent so almost no ions entering the HRIM module are wasted (∼100% ion utilization efficiency). The system can also be operated in a passthrough mode where ions are not accumulated and instead directly pass through the HRIM module losslessly without ion mobility filtering, enabling standard UHPLC-HRMS analysis to be performed.

**Figure 1:**
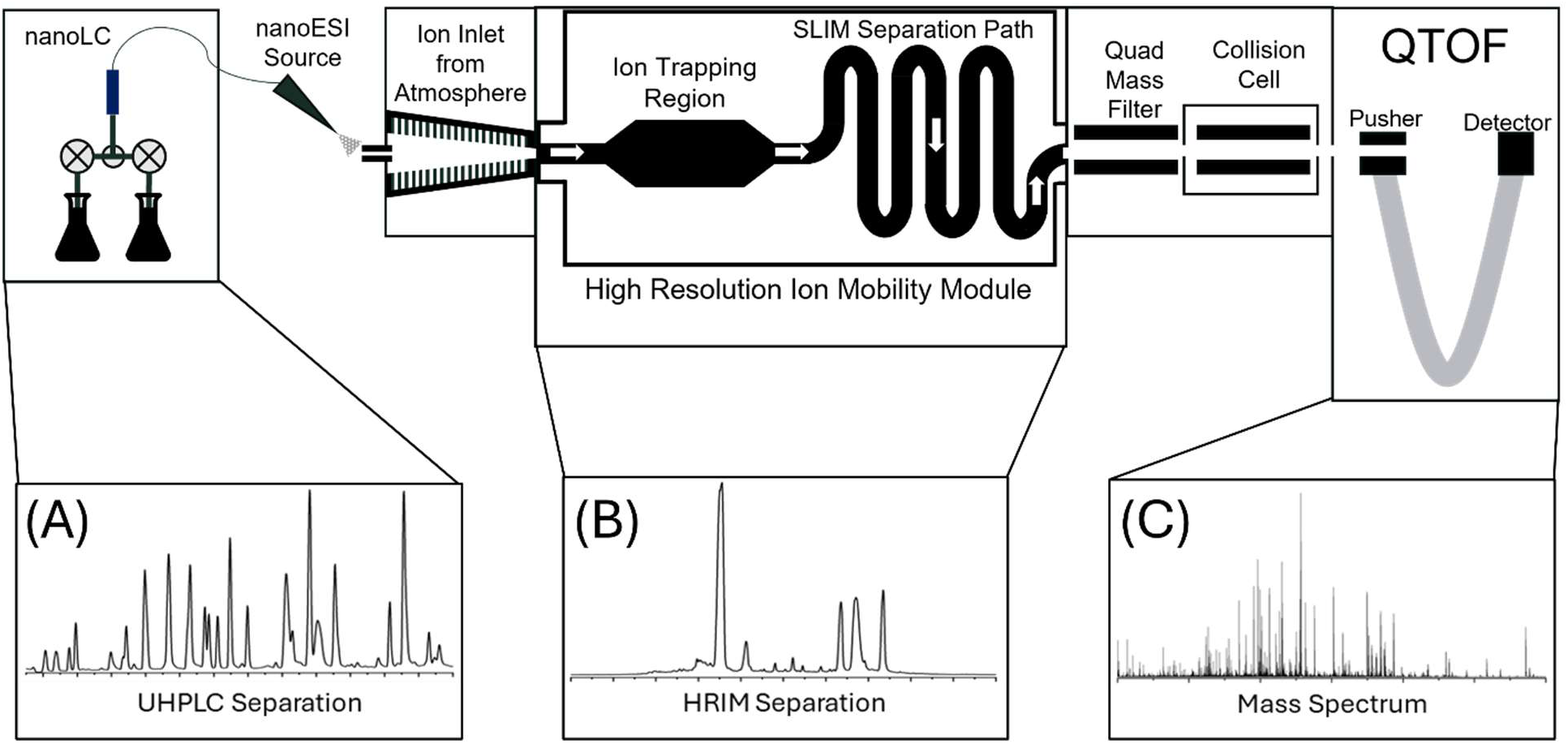
Schematic overview of the HRIM/QTOF system configured for PAMAF mode testing. Samples can be introduced using UHPLC generating chromatographic separation (**A**). Samples are ionized by nano-electrospray ionization and introduced into the HRIM module where they undergo high resolution ion mobility separation via SLIM (**B**). After separation, ions transit into the QTOF region where they undergo mass analysis or CID followed by mass analysis (**C**).

After passing through the HRIM system, ions enter the QTOF mass analyzer (**Fig. 1C**) where they undergo mass analysis with no collision energy applied to generate full-scan mass spectra (MS1) or mass analysis following application of collision energy to generate fragment ion mass spectra (MS2). The system was configured to allow MS1 and MS2 events to happen back-to-back, generating alternating MS1 and MS2 frames of data. In certain scenarios, the system was also configured to generate multiple MS2 frames of data between MS1 frames (discussed below).

Since HRIM separation occurs prior to ion fragmentation, subsequent ion mobility separation events (i.e. back-to-back MS1 and MS2 frames) are always the separation of unfragmented precursor ions. This allows for intuitive association of fragment ions from each MS2 frame where collision energy was applied to their corresponding precursor ion identity in a neighboring MS1 frame where collision energy was not applied. This mobility-aligned fragmentation (MAF) methodology is highlighted in **Figure 2**. **Figure 2A** displays the extracted ion chromatogram (XICs) for an example precursor ion from a HeLa digest sample ([M+2H]^2+^, ISGLIYEETR) and its dominant fragments. Each point in the XICs represents either an MS1 (green) or MS2 (red) ion mobility frame composed of both mobility and associated mass spectra occurring at each point in the mobility separation. This produces ion mobility vs. *m/*z heatmaps as shown in **Figures 2C** and **2E**, which correspond to adjacent MS1 and MS2 frames, as shown in **Figure S2**. **Figures 2B** and **2D** show to the summed mass spectra from each ion mobility separation frame. The heatmap shown in **Figure 2F** highlights the unique capabilities of mobility alignment, where precursor #3 from **Figure 2C** (green) is aligned with the fragments (red) shown in **Figure 2E**. This alignment of the fragments with the precursor allows for mobility filtering of the fragmentation mass spectrum, shown in **Figure 2G**. By taking “slices” of the mobility dimension in both MS1 and MS2 frames at the same arrival time, the fragmentation spectrum can more easily be annotated and matched to a peptide sequence for positive identification.

**Figure 2:**
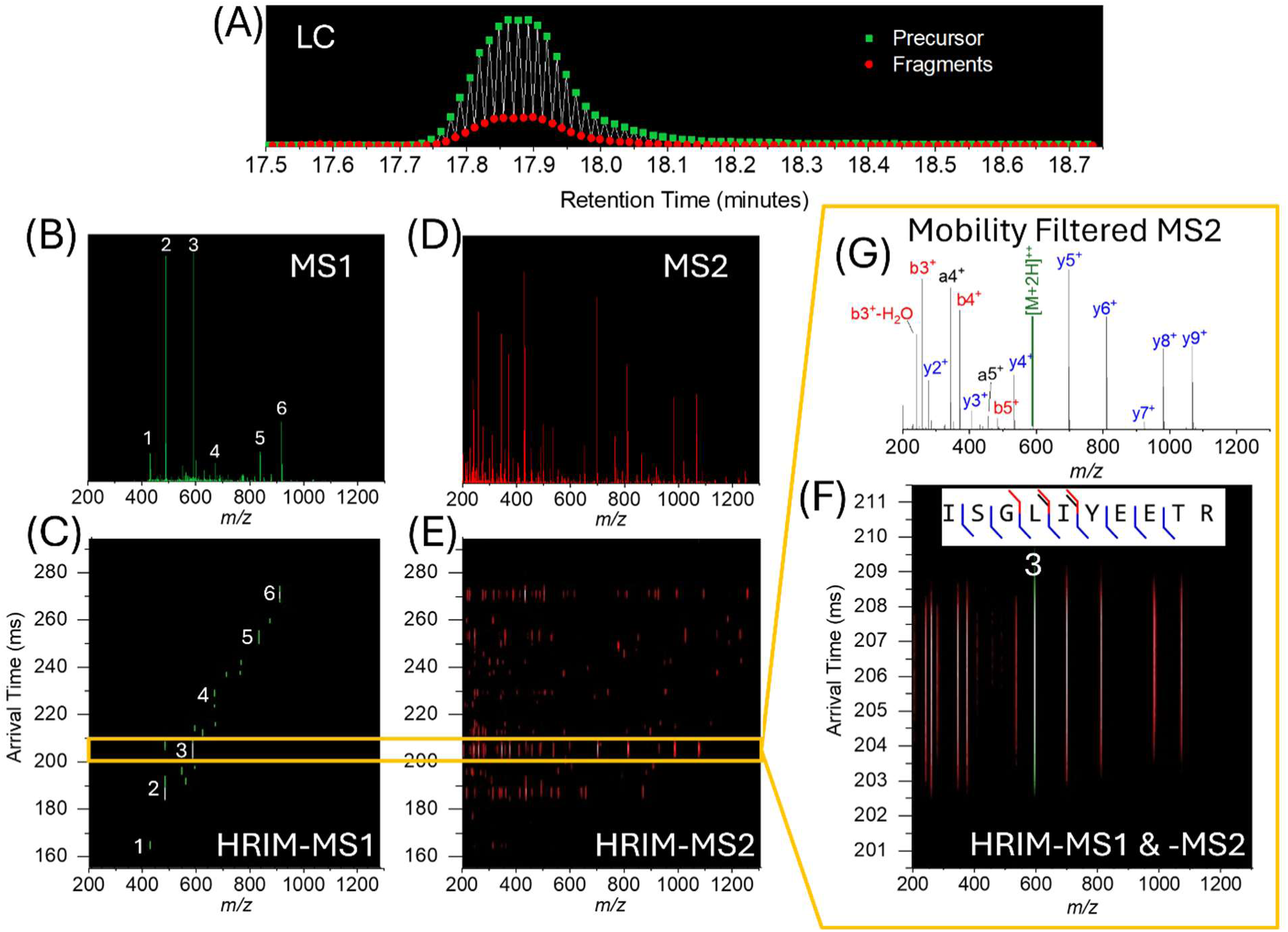
(**A**) In a PAMAF analysis, alternating MS1 (precursor) and MS2 (fragment) scans are acquired. Each MS1 scan is comprised of a total mass spectrum (**B**) that can be further expanded into a mobility vs. m/z frame (**C**). Each MS2 scan contains all of the fragments in that scan (**D**) and can also be expanded into a mobility vs. m/z frame (**E**). With MAF, precursors from the MS1 frame can be aligned to fragments observed in the MS2 frame (**F**). MAF reduces complexity of the fragmentation spectrum allowing for higher confidence identifications (**G**).

Figure 3 provides another example of how MAF can be used to attribute fragment ions to their precursor. In the top portion of Figure 3A depicts the extracted ion mobilograms (XIMs) for precursor ions ([M+2H]^2+^) of the 15 peptides present in the PRTC mixture (Precursor *m/*z, Fig. 3B). The bottom portion of this figure shows the XIMs for the major fragment ion for each of the 15 peptides (Fragment *m/z*, Fig. 3B).

**Figure 3:**
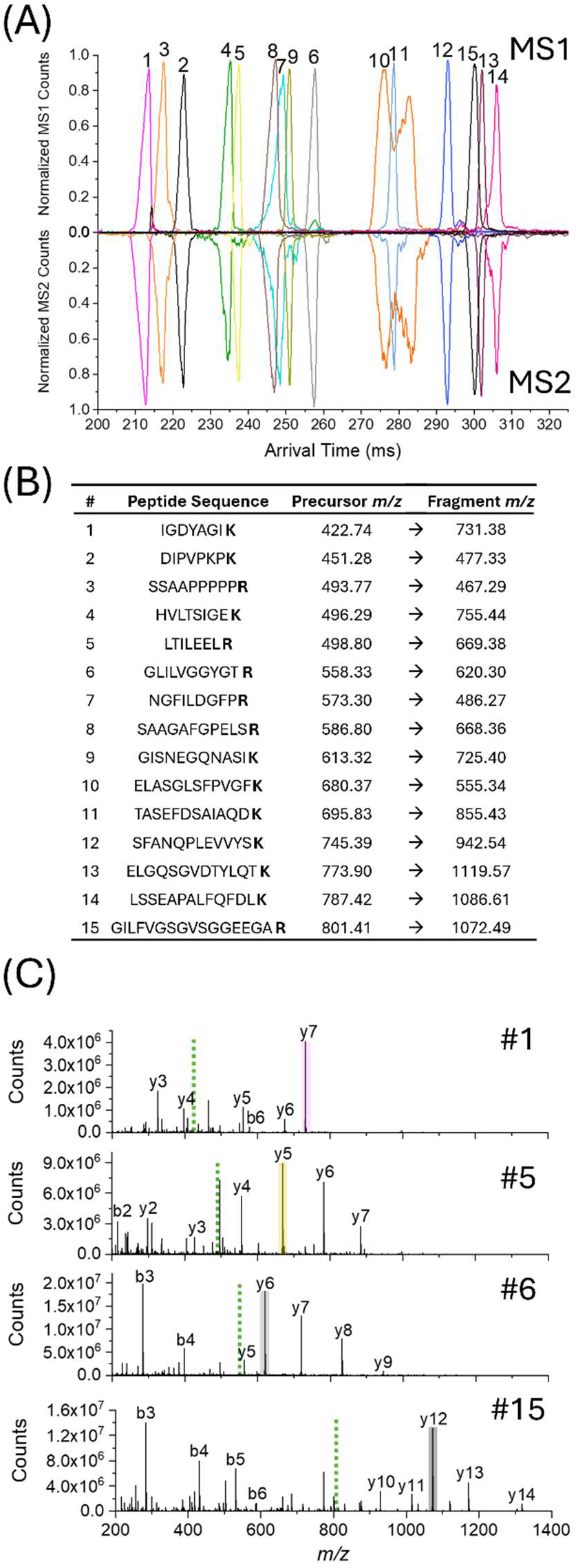
(**A**) Comparative alignment of the XIMs for the precursor ions of the 15 peptides in PRTC (top plot) to the XIMs for a selected diagnostic fragment (bottom plot) of each of the 15 peptides highlighting the mobility alignment of fragments to precursors in MAF analyses. (**B**) Table showing the peptide sequence, precursor m/z and fragment m/z for the XIMs plotted in A. (**C**) Selected MS2 fragmentation mass spectra for 4 of the peptides. Fragments plotted in A are shaded and precursor position shown by green dotted line. In MAF, every point in the mobility dimension produces a mass spectrum which can be used to mobility align fragments with a specific precursor.

The top and bottom XIMs are representatives of consecutive MS1 and MS2 frames. It should be noted that the fragment XIMs are aligned with their precursor XIMs, and the mobility peak shapes are conserved. Figure 3C shows examples of the full MS2 spectra that were acquired for four of the 15 peptides. In these spectra, the peak used for the MS2 XIM in Figure 3A is highlighted as well as the position of the precursor (green dashed line). In MAF, a full mass spectrum is collected for every point in the mobility separation, meaning that every fragmentation spectrum can be potentially matched back to a precursor in the previous MS1 frame.

As discussed above, many modern HRMS systems rely on quadrupole filtering for enhancing the fragmentation spectral performance by lowering the complexity of the fragmentation spectrum by allowing only precursors within a narrow range of *m/z* values to be fragmented per MS2 spectrum. Operating in DIA mode with a narrow window of *m/z*, many MS2 spectra must be acquired per cycle to cover the entire mass range of interest. To keep cycle times down (as defined by the time between MS1/precursor scans), MS2 dwell times must be correspondingly short, with systems often operating with dwell times as low as 5 ms, or 200 Hz^36^. Utilizing short dwell times may mean that less than 0.5% of the available ions for a particular precursor are ultimately transmitted, fragmented, and detected. Conversely, by accumulating ions in parallel with HRIM separation and not relying on quadrupole filtering, virtually all ions can be transmitted for fragmentation. Figure 4 compares how the signal level for an infusion of 50 fmol/µL of the PRTC mixture differs between a quadrupole filtering experiment (without HRIM separation) operating at a 200 Hz MS/MS rate and data collected in PAMAF mode capable of an effective MS2 rate exceeding 500 Hz. In Figure 4A, the red trace depicts the XIM for the precursor ion of GILFVGSGVSGGEEGA**R** ([M+2H]^2+^, *m/z* 801.41) after undergoing HRIM separation. The blue trace represents the same signal without HRIM separation (passthrough mode) over the same time period. Both traces are the sum of 6 seconds worth of signal, representing the baseline width of a typical UHPLC peak^37^. In PAMAF mode, the signal for this peptide is focused into a narrow peak (∼2 ms FWHM) as compared to a continuous, lower-level signal generated in passthrough mode. Figure 4B shows the MS2 signal that would be present with a 5 ms dwell time (200 Hz) MS2 experiment with in passthrough mode (Fig. 4B top, highlighted area from blue trace in **Fig 4A**) as compared to the MS2 signal resulting from 2 ms of PAMAF mode data (Fig. 4B bottom, highlighted area from red trace in **Fig 4A**). Even with 2.5x less dwell time, the PAMAF data has ∼150x higher signal, demonstrating the effect of ion utilization on the resulting MS2 sensitivity. In terms of MS2 rate, generating MS2 spectra from PAMAF mode data every 2 ms corresponds to an effective rate of 500 Hz, with mobility peak deconvolution enabling even higher rates.

**Figure 4:**
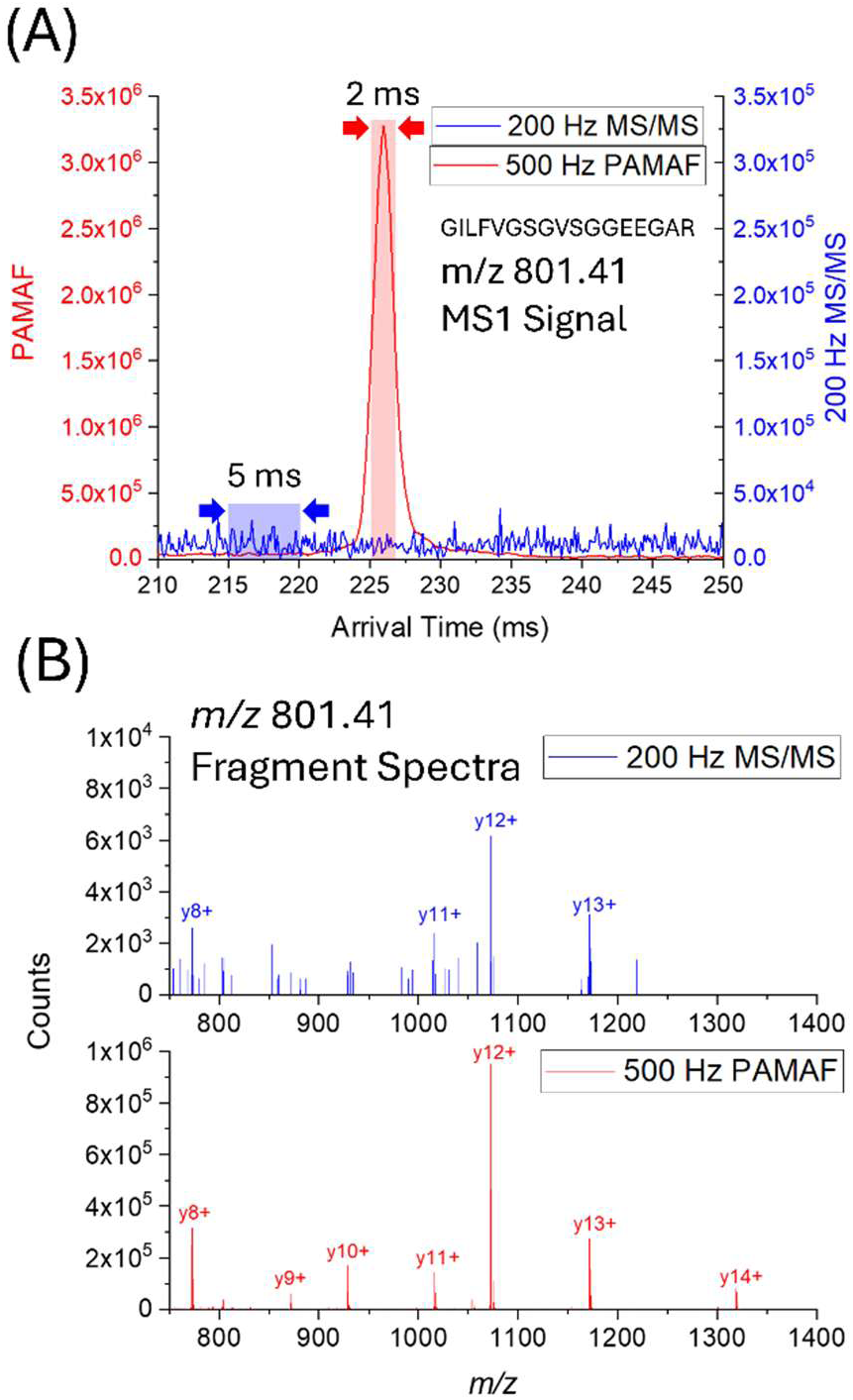
(**A**) Comparison XIM signal for precursor ion of GILFVGSGVSGGEEGAR (m/z 801.41) from PRTC in passthrough mode with no mobility filtering (blue trace) to HRIM separation of the same ion (red trace) (**B**) Top spectrum (blue trace) represents a 5 ms integration of MS2 signal for passthrough mode compared to the bottom spectrum (black trace) which represents a 2 ms integration of MS2 signal for HRIM data.

### Benchmarking

To evaluate the performance of PAMAF mode, benchmarking experiments were performed using HeLa protein digest in two workflows: low-load (0.1-10 ng) and high load (25-1000 ng) for global shotgun proteomics. As a baseline, DDA analysis was first conducted using a standard Agilent QTOF workflow^38^, operating the MOBIE system in passthrough mode as described above. For low-load proteomics, including single cell equivalent inputs, performance was assessed at varying sample speeds (Figure 5A). At 40 SPD, DDA detected no proteins below 0.5 ng which equates to approximately 3 processed HeLa cells^39^ and identified only 238 protein groups (PGs) at 10 ng. However, PAMAF consistently produced higher identifications at low loads: 296, 1409, and 2356 PGs at 0.1, 0.2, and 0.5 ng, respectively, increasing to 3206 PGs at 10 ng. At 80 SPD, PAMAF outperformed 40 SPD for sub-nanogram inputs that yield 1028, 1707, 2538, and 2907 PGs at 0.1, 0.2, 0.5, and 1 ng, respectively. We hypothesize increased performance for the 80 SPD method is attributed to a reduction in LC peak width which increased the signal level per frame. At 120 SPD, the number of identifications was lower than at 80 SPD, likely due to a reduction in LC resolution. Protein groups quantified at CV < 20% and < 10% in PAMAF mode were similar across method speeds at the same sample load (**Figure S5A**). Overall, these results demonstrate that the increased ion utilization efficiency of the PAMAF operation mode results in a substantial increase in detection sensitivity, which is the primary performance challenge for sample limited analyses.

**Figure 5:**
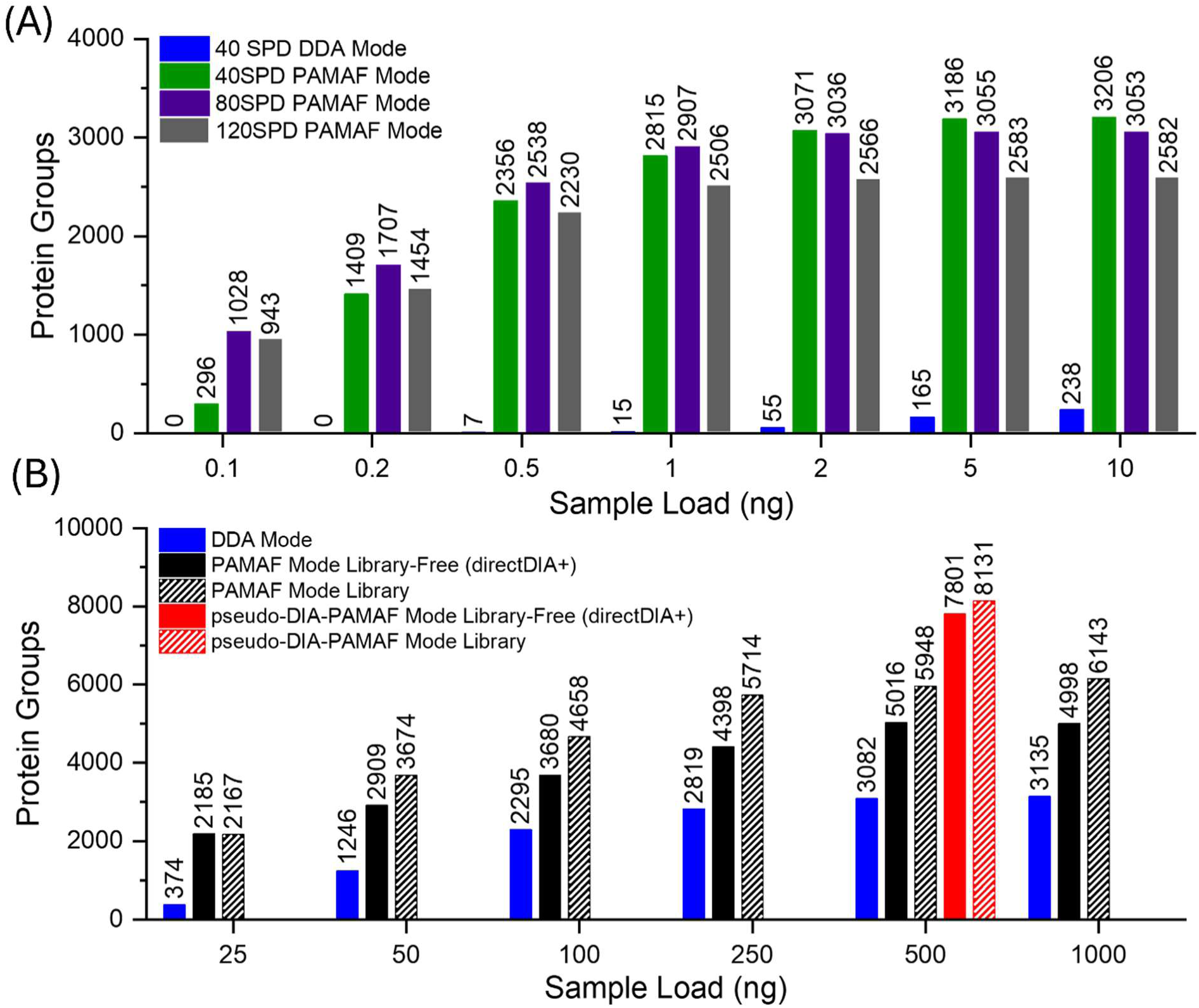
**(A)I**dentified protein groups from HeLa digests (100 pg - 10 ng) acquired in DDA and PAMAF modes at sample speeds of 40 SPD, 80 SPD, and 120 SPD. Data were processed in Spectronaut 19 using library-free (directDIA+) search; (**B**) Identified protein groups obtained from HeLa digests (25 – 1000 ng) using DDA, PAMAF, and pseudo-DIA-PAMAF modes at same sample speed of 30 SPD. Data were processed in Spectronaut 19 using two distinct search strategies: library-free (directDIA+) and library-based search.

Similar benchmarking of PAMAF versus DDA acquisition was performed for higher load levels of HeLa digest (25-1000 ng) at 30 SPD on the same EvoSep One system with identical columns and emitters (Figure 5B). In DDA mode, PG identifications increased from 374 at 25 ng to 3135 at 1000 ng, with identification rates plateauing around 500 ng. PAMAF mode delivered a marked improvement, rising from 2185 at 25 ng to 4998 at 1000 ng using a library-free (directDIA+™ HeLa FASTA) search in Spectronaut v19, reflecting improved sensitivity and specificity. Further enhancement was achieved using a spectral library generated from 24 fractionated HeLa samples analyzed in PAMAF mode (**Fig S1, Table S1**), which yielded PG IDs from 2167 at 25ng to 6143 at 1000 ng.

This represents a 23% increase at higher loads compared to the results from the library-free search. Quantitative precision was also improved in PAMAF compared with DDA (**Fig. S5B**) as determined from values below %CV thresholds, as described above.

While the exclusive use of HRIM isolation in PAMAF mode provides levels of specificity on par with DIA quadrupole isolation, previous work has demonstrated improved precursor specificity can be achieved by combining HRIM separation with quadruple *m/z* isolation^40^. Due to the limited software integration between the MOBIE and QTOF systems, dynamic adjustment of the quadrupole isolation during mobility separation as performed in the case of DIA-PASEF^19^ was not possible on the current prototypes.

However, in an effort to assess the effect of combined HRIM and quadrupole isolation in a proof of concept experiment that splits various static quadrupole isolation events into replicate sample injections, a pseudo-DIA-PAMAF mode of operation was explored (**Fig. S3** and **S4)**. This approach is conceptually similar to gas phase fractionation (GPF) on FAIMS-Orbitrap systems^41,42^. In pseudo-DIA-PAMAF mode, HRIM separation and quadrupole *m/z* isolation were applied simultaneously for all MS2 scans, while MS1 scans were acquired without quadrupole isolation. A total of 20 injections were performed spanning the *m/z* range of 390-1190 Th. Each injection employed two constant quadrupole isolation windows (20 Th each), with a sequence of one MS1 scan followed by two MS2 scans. For each MS2 scan, a single isolation window was used, yielding 40 total windows to cover the entire precursor *m/z* range (2 windows per injection x 20 injections). The cycle time of one MS1 scan (445 ms) and two MS2 scans (440 ms each) was about 1.325 s. All MS1 and MS2 frames used the same on-board accumulator (OBA) fill time of 390 ms. All other parameters were identical between the PAMAF and pseudo-DIA-PAMAF modes, with collision energies (CEs) optimized individually.

Figure 5B compares the number of PG identifications in DDA, PAMAF, and pseudo-DIA-PAMAF modes, with PGs quantified at < 20% and < 10% CVs shown in **Figure S5B**. Pseudo-DIA-PAMAF was tested at 500ng HeLa, achieving 56% more PGs in library-free searches and 37% more with the library search compared to PAMAF mode results. This improvement demonstrates how adding quadrupole filtering to MAF further increases the specificity of precursor isolation, enabling cleaner MS2 spectra leading to enhanced peptide identification^40^. Assuming this pseudo-DIA-PAMAF workflow could be translated to a fully automated acquisition on a single injection, the detection of 8,131 PGs represents performance on par with other leading-edge proteomics platforms^10,43–47^.

### Quantitative Performance Assessment

To test quantitative performance, the MS Qual/Quant QC mix was analyzed at 200 SPD in both DDA and PAMAF modes. The MS Qual/Quant QC Mix contains six, trypsin-digested proteins with two to three heavy stable isotope labeled peptides (C-terminus, +8 Arginine and +10 for Lysine) per protein at various abundance ratios (**Table 1**, CoA Measured ratio). The sample was diluted such that final tested amounts for the heavy labeled peptides ranged from 50 fmol to 0.04 fmol on column. Both sets of data were processed using Skyline with MSAmanda^48^, leveraging features identified by Spectronaut in the case of PAMAF data.

**Table 1:**
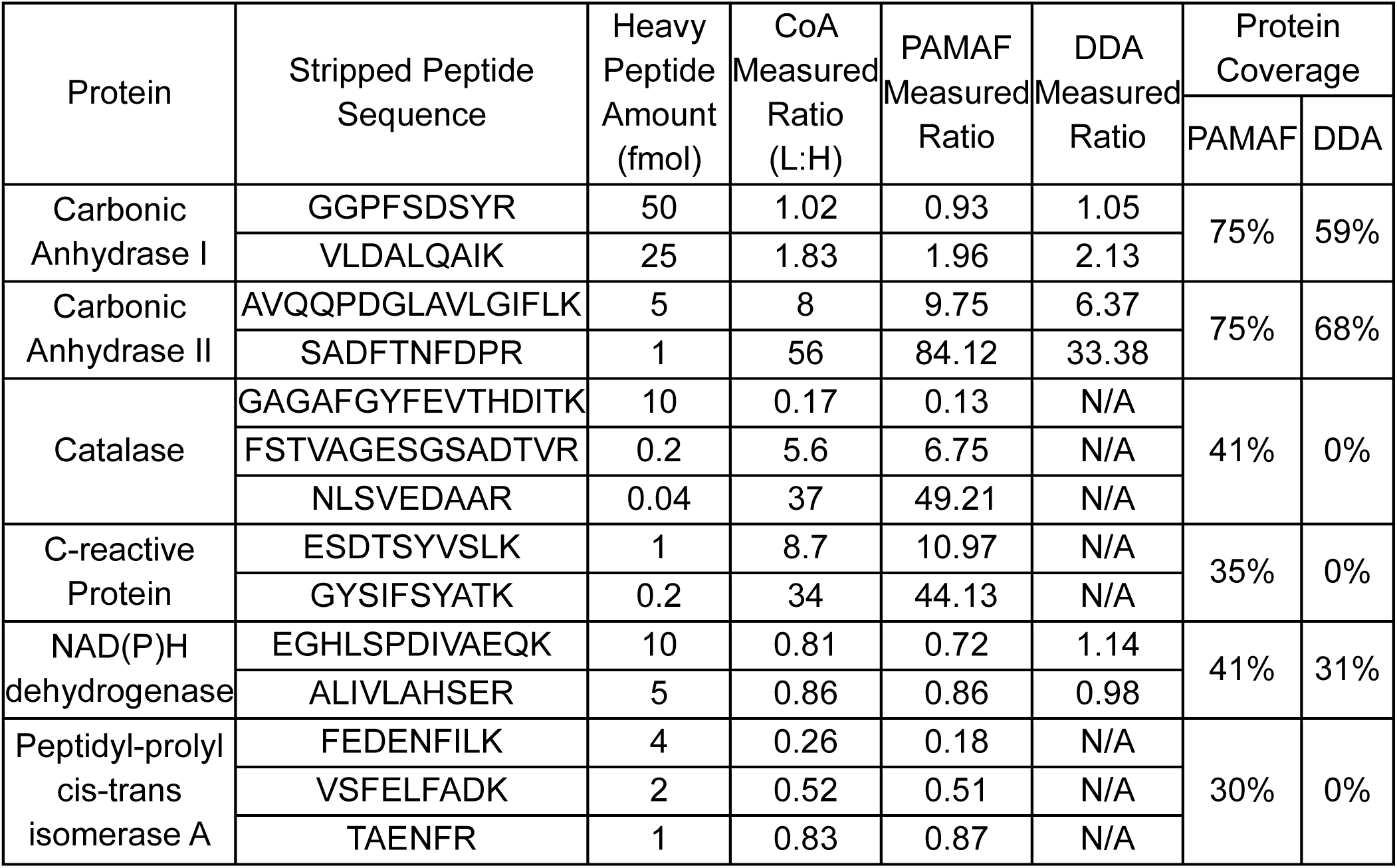
Outline of proteins present in the Qual/Quant mixture. For each protein 2-3 heavy labeled peptides (+8 for K and +10 for R) are spiked in at various ratios. Certificate of analysis values for those ratios are listed below. PAMAF mode and DDA mode (where detected) measured ratios are shown along with percent protein sequence coverage for each mode.

Figure 6A is a visualization of HRIM-MS data for the Light: Heavy labeled pair of ions corresponding to the [M+2H]^2+^ of the peptide GAGAFGYFEVTHDITK showing the expected 0.2:1 abundance ratio with intuitive alignment of both precursor ions in the HRIM dimension. Figure 6B details the expected and measured Light:Heavy ratios of all peptides in the standard mixture. Using Spectronaut’s internally calculated MS2-based Heavy to Light ratio and Skyline’s reported MS1 values, all PAMAF mode Light:Heavy ratios were measured to be within the manufacturer’s tolerance specified per peptide. Passthrough DDA mode values were calculated from a Skyline/MSAmanda search where, in some cases, one or both target peptides were not detected. In general, peptides below 5 fmol were below the limit of detection in DDA mode. However, with PAMAF mode, quantitation of all listed Light:Heavy peptides were possible and provided higher quantitative accuracy than DDA mode on average.

**Figure 6:**
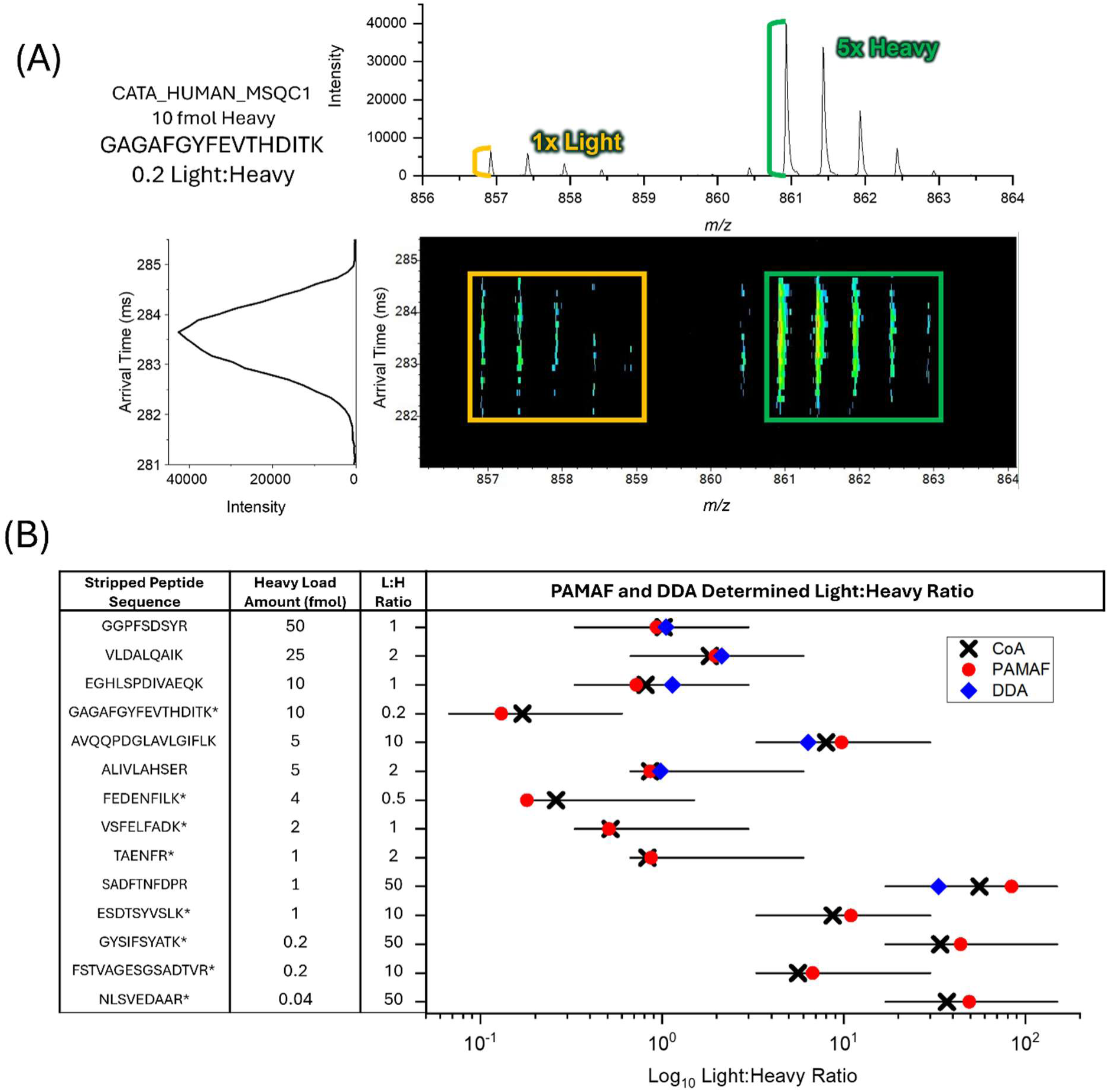
**(A)** Visualization of the MS1 mobilogram, mass spectrum and mobility/mass heat map for one of the light:heavy pairs (GAGAFGYFEVTHDITK) in the Qual/Quant Mixture. The heavy peptide is present at 5x the light level (10 fmol vs. 2 fmol). (**B**) Detailed plot of the expected light to heavy ratio for each of the peptide pairs in the mixture. Plot shows the accepted sample range (black bar), certified sample value (CoA, black X), PAMAF value (red dot), and DDA value, if present (blue diamond). Peptides with an asterisk (*) were not identified/quantified in DDA.

MS Qual/Quant QC Mix analyzed in PAMAF mode demonstrated high accuracy relative quantitation over a relative abundance range of 0.2:1 to 1:50. The peptide with the lowest sample load at 0.04 fmol (NLSVEDAAR) was detected with an average L:H ratio error of ∼2% compared to the theoretical ratio (Fig. 6B). This peptide was not detected via DDA analysis in either form. Average values for PAMAF mode as low as 0.46% difference from the certificate of analysis (CoA) measured ratio were achieved for a more abundant peptide (ALIVLAHSER at 5 fmol), with the respective DDA difference at 13.45% (**Table 1**). PAMAF mode detected and quantified peptides covering >3 orders of magnitude concentration range (0.04 fmol to 50 fmol). In addition to PAMAF mode also detecting and quantifying all 14 peptide pairs, protein sequence coverage was higher as compared to DDA mode, which did not identify several of the proteins.

### Isobars and Isomer Separation

The value of PAMAF mode operation extends beyond the speed and sensitivity improvements described in previous sections. The reliance of conventional MS/MS operation on quadrupole filtering to isolate ions for fragmentation renders the analysis blind to most isomeric and isobaric interferences that are present in complex samples^49^. These compounds are largely ignored or lead to low-quality spectral matches that lower identification confidence for the convoluted spectra. PAMAF mode overcomes this challenge of overlapping isobaric and isomeric precursors that lead to chimeric fragmentation spectra. Figure 7 shows two examples of coeluting isobaric (Fig. 7A**-1**) and isomeric (Fig. 7B**-1**) peptides that can be physically resolved by HRIM separation prior to fragmentation, enabling extraction of clean fragmentation spectra. For Figure 7A, a pair of coeluting isobaric peptides (AIDEVNR, *m/z* 408.7141 and EIQTAVR, *m/*z 408.7323) present in the HeLa digest standard analyzed at 30 samples per day are shown. Figures 7A**-2** and **A-3** show the HRIM-MS1 heatmap and mobilogram for the two peptides (#5 and 6) showing separation of the two isobars. With quadrupole only filtering (including narrow band isolation), both isobaric peptides would be transmitted simultaneously, producing a chimeric fragmentation spectrum (Fig. 7A**-4**) comprised of fragments from both peptides. Accurate identification of either peptide from this combined spectrum is unlikely, particularly when one peptide is in higher abundance than the other. However, with PAMAF-mode, MAF separates these two isobaric peptides, producing higher quality fragmentation spectra (**Fig 7A-5** and **A-6**) which can be more readily matched to a sequence.

**Figure 7:**
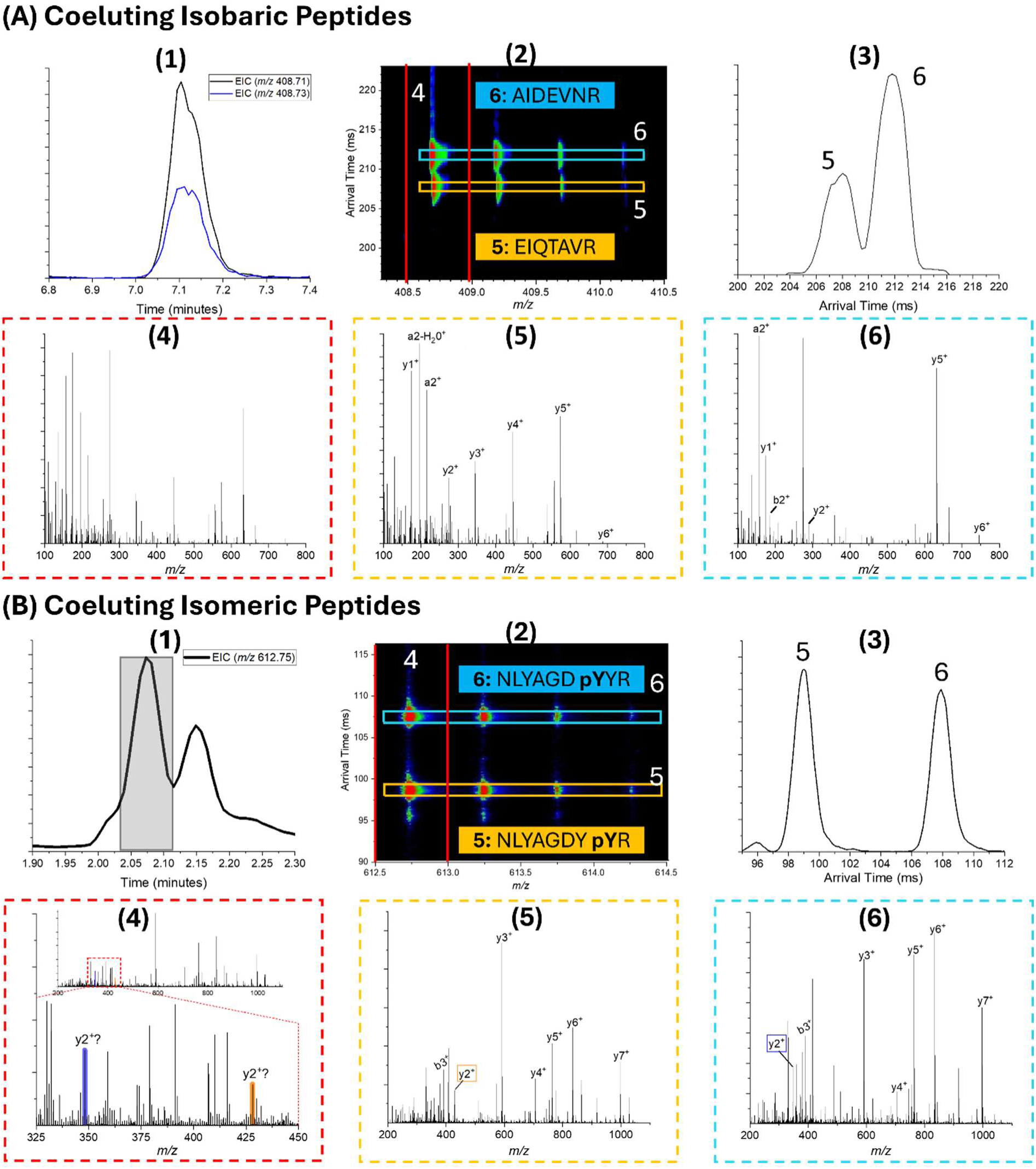
**(A)(1)** XIC for coeluting isobaric peptides AIDEVNR (m/z 408.7141) and EIQTAVR (m/z 408.7323). (**2**) Arrival Time vs. m/z heatmap and (**3**) XIM of the two peptides showing separation. (**4**) Chimeric fragmentation spectrum corresponding to isolation based on monoisotopic m/z vs. (**5/6**) clean fragmentation spectra based on mobility precursor selection. **(B)(1)**XIC for isomeric peptides (m/z 612.7475) with differing localization of phosphorlyation. The highlighted region contains two coeluting isomeric peptides shown in **(B)(2,3)**. **(2)** Arrival Time vs. m/z heatmap and **(3)** XIM of the two peptides showing separation. **(4)** Chimeric fragmentation spectrum corresponding to isolation based on monoisotopic m/z vs. **(5/6)** clean fragmentation spectra based on mobility precursor selection.

Another example shown in Figure 7B details a series of isomeric peptides from the phosphorylated peptide standard analyzed at 200 samples per day. These isomers of NLYAGDYYR only differ in the location of the phosphorylation post-translational modification (PTM) of the tyrosine (Y) residue, with three potential sites available. Figure 7B**-1** shows the extracted ion chromatogram for the [M+2H]^2+^ precursor ion of the phosphorylated peptide. In the chromatogram, there are two resolved peaks, indicating at least two species are present. By using mobility to examine the first chromatographic peak further (highlighted region in Fig. 7B**-1**), two isomeric species were observed in the mobility domain. Figures 7B**-2** and **B-3** show the HRIM-MS1 heatmap and mobilogram for the highlighted region, showing two mobility resolved peptides (7B-5 and 7B-6). With quadrupole only filtering (including narrow band isolation), both isomeric peptides would be transmitted, producing a chimeric fragmentation spectrum (Fig. 7B**-4, inset**). With PAMAF mode, MAF analysis separates these two isomeric peptides, producing higher quality fragmentation spectra (**Fig 7B-5** and **B-6**) which enable accurate identification of these peptides that differ only by PTM location. The key fragment ion for phosphate localization is the y2 fragment ion. Since the position of the phosphate in the isomeric peptides differs by only one residue, identifying the y2 fragment confirms the presence (y2 of peptide 7B-5) or absence (y2 of peptide 7B-6) of the phosphate on the tyrosine residue directly adjacent to the peptide’s C-terminus. In the quadrupole only isolation, both y2 fragments are observed, indicating a chimeric fragmentation spectrum that would inhibit typical search programs from confidently assigning the PTM to the Y7 versus Y8 position.

## DISCUSSION

Here we described and evaluated the novel PAMAF mode as an alternative approach to operation for quadrupole time-of-flight mass analysis leveraging high resolution ion mobility in place of, or in addition to, quadrupole precursor isolation. This methodology is particularly well suited to the analysis of complex proteomics samples where high sensitivity data-independent acquisition of LC-MS data is desired. PAMAF mode operation provided high-speed MS2 spectral generation (>500 Hz) without sacrificing sensitivity of detection, which is a typical tradeoff in quadrupole-based analysis. The key principle of the PAMAF mode is MAF, where precursor fragmentation occurs after mobility separation. This allows for alternating MS1 and MS2 mobility frames of data to be collected and fragments observed in the MS2 frame to be associated back to precursors observed in the MS1 frame based on their conserved mobility peak position. MAF provides specificity of precursor/fragment pairs without throwing away ion signal via quadrupole filtration. Additionally, the use of parallel accumulation allowed for all ions entering the HRIM module from the ionization source to be utilized for analysis.

Benchmarking studies showed that compared to baseline DDA analysis, PAMAF mode offered much higher protein IDs with greater confidence in the quality of those IDs. In cases where analysis is specificity limited, the addition of quadrupole filtering to produce pseudo-DIA-PAMAF mode data showed further improvements in protein detection. Quantitative performance of the PAMAF mode was better than the baseline DDA mode in all cases. The high resolving power of PAMAF mode can also be leveraged to reduce chimeric overlap in fragmentation spectra, greatly improving detection of coeluting isomers and isobars, with particular applicability to the challenge of PTM site localization.

We believe the PAMAF operation mode represents an important addition to the field of bottom-up proteomics analysis with significant potential to extend the performance profile of future LC-MS systems.

This article contains supplemental Figures and Tables.

## Abbreviations

AT: Arrival time
CCS: Collision cross section
CE: Collision energy used for MS/MS fragmentation by CID
CID: Collision-induced dissociation
CoA: Certificate of Analysis
DDA: Data-dependent acquisition
DDM: n-Dodecyl β-D-maltoside
DIA: Data-independent acquisition
DNIF: Dual narrow ion funnel
FDR: False discovery rate
HDMS^E^: High-definition mass spectrometry
HRIM: High-resolution ion mobility
HRMS: High-resolution mass spectrometry
iRT: Indexed retention time
MS: Mass spectrometry
MS/MS: Tandem mass spectrometry
MS1: Precursor mass spectrum scan
MS2: Fragmentation mass spectrum scan
MAF: Mobility aligned fragmentation
OBA: On-board Accumulator
PA: Parallel accumulation
PAMAF: Parallel Accumulation-Mobility Aligned Fragmentation
PASEF: Parallel Accumulation Serial Fragmentation
PCB: Printed circuit board
PG: Protein group
PRTC: Peptide retention time calibration
PTM: Post translational modification
Q-TOF: Quadrupole Time-of-Flight mass spectrometer
R_p_: Resolving power
SIL: Stable isotope labeled
SPD: Samples per day
SLIM: Structures for Lossless Ion Manipulation
Th: Thomson
TIMS: Trapped ion mobility spectrometry
TW: Traveling wave
TWIMS: Traveling wave ion mobility spectrometry
UDMS^E^: Ultra definition mass spectrometry
UHPLC: Ultra-high performance liquid chromatography
XIC: Extracted ion chromatogram
XIM: Extracted ion mobilogram

