## Supplemental Figures and Tables for "A Parallel Accumulation-Mobility Aligned Fragmentation Strategy Utilizing High-Resolution Ion Mobility for High Performance Proteomics Analysis"

Leonard C. Rorrer, III[1*], Liulin Deng[1], Lauren Royer[1], Isabel Uribe[1], Benjamin C. Orsburn[2], Oliver Bernhardt[3], Tejas Gandhi[3], Lukas Reiter[3], Daniel DeBord[1]

*Corresponding Author

*Author affiliations;*
*1: MOBILion Systems, Inc. 4 Hillman Drive, Suite 130, Chadds Ford, PA  19317, United States*
*2: Organ Pathobiology and Therapeutics Institute, University of Pittsburgh, 3501 5$^{th}$ Avenue, Pittsburgh, PA  15260, United States*
*3: Biognosys, Wagistrasse 21, 8952 Schlieren, Switzerland*

SUPPLEMENTAL INFORMATION CONTENTS


| | |
|---|---|
| Sample Fractionation for Library Development | S-3 |
| Figure S-1: Protein Groups Identified in Fractions | S-4 |
| Table S-1: Fractionated Library Details | S-4 |
| Figure S-2: Schematic of PAMAF Mode Workflow | S-5 |
| Figure S-3: Schematic of pseudo-DIA-PAMAF Mode Workflow | S-6 |
| Figure S-4: Overview of Quadrupoles Windows in pseudo-DIA-PAMAF Mode | S-7 |
| Table S-2: pseudo-DIA-PAMAF Mode Parameters | S-7 |
| Figure S-5: Quantifiable Protein Groups for Low and High Load Benchmarking | S-8 |

*Sample Fractionation for Library Development:* A set of fractionated HeLa digest samples were collected to use for generating a reference processing library for comparison of system benchmarking data. The 100 µg of the HeLa proteomic standard (described above) was resuspended in high pH fractionation buffer A (4.5 mM ammonium formate, 2% acetonitrile) and loaded on a 250 mm x 4.6 mm Zorbax extend-C18 Rapid Resolution column (3.5 µm beads, Agilent 763953-902). The samples were eluted with a 109 minute gradient of mobile reagent B (4.5 mM ammonium acetate, 90% acetonitrile) at a flow rate of 0.8 mL/min. The gradient was 0% B for 13 minutes, rising to 16% B for 60 minutes, 40% B for 40 minutes, 44% B for 5 minutes, 60% B for 13 minutes, 99% B for 4 minutes and dropping back to 0% B for the remaining 10 minutes. Forty-eight fractions were eluted, collected, and then concatenated down to 24 in the following order: 1 and 25, 2 and 26, 3 and 27, 4 and 28, etc. Each dried sample had approximately 3 µg of material based on absorbance and based on this, samples were diluted to approximately 10 ng/µL in water with 0.1% formic acid (v/v) prior to analysis by UHPLC.

To generate a fractionated library for use in data processing, each fractionated sample was analyzed by UHPLC/HRIM/Q-TOF PAMAF mode at 30 SPD (3 replicates at ~200 ng loading per fraction). The results were processed in Spectronaut 19 in a library-free mode (directDIA+). **Supplemental Figure 1** shows the number of identified protein groups for each fraction. The processing results were combined using the Spectronaut 19 standard procedure for generating libraries from results. **Supplemental Table 1** shows the details of the library output from this process. This resulting library was used for library (DIA Analysis) processing in Spectronaut.

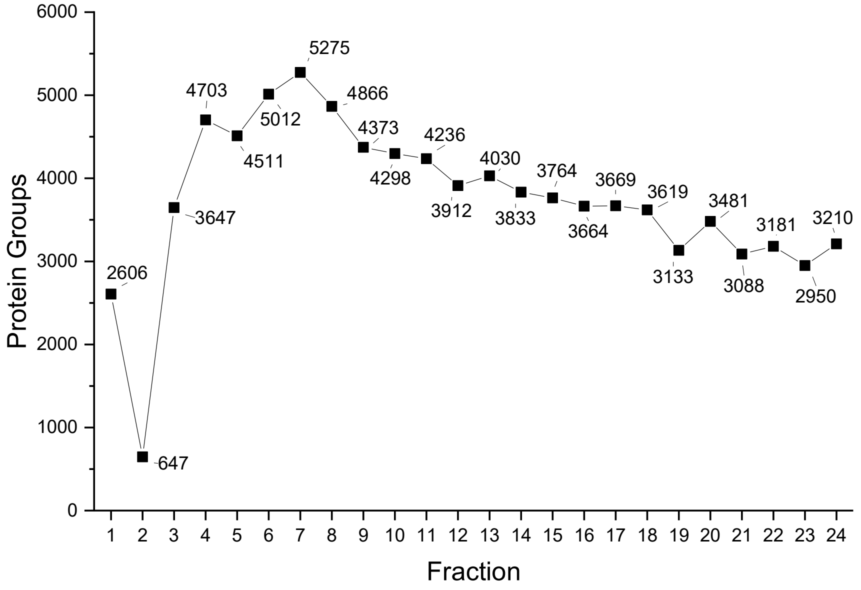


**Figure S1**: Protein groups identified for each fraction collected from the fractionated HeLa sample. Processing was done on each fraction sample using Spectronaut 19 in library-free mode (Direct DIA+).

| **Fractionated Library Details** | |
|---|---|
| Protein Groups | 10,316 |
| Precursors | 184,109 |
| Modified Peptides | 141,172 |
| Peptides | 126,407 |

**Table S1**: Details of the library generated from the fractionated HeLa samples. Library created from combining processing results of 24 fractions shown in Supplemental Figure 1.

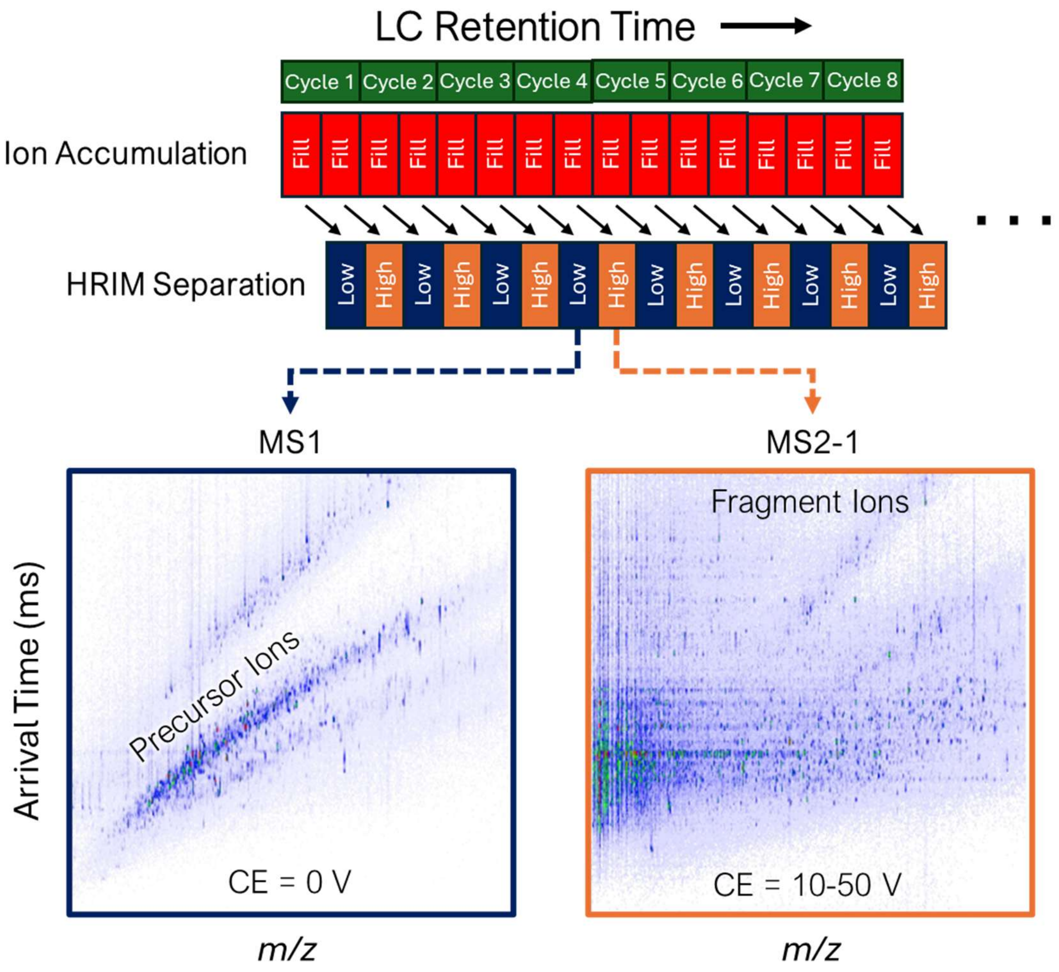


**Figure S2**: Schematic diagrams of the PAMAF mode workflow. In PAMAF mode, alternating low CE (MS1) frames and high CE (MS2) frames are collected. Using MAF, fragment ions are matched to precursors for identification.

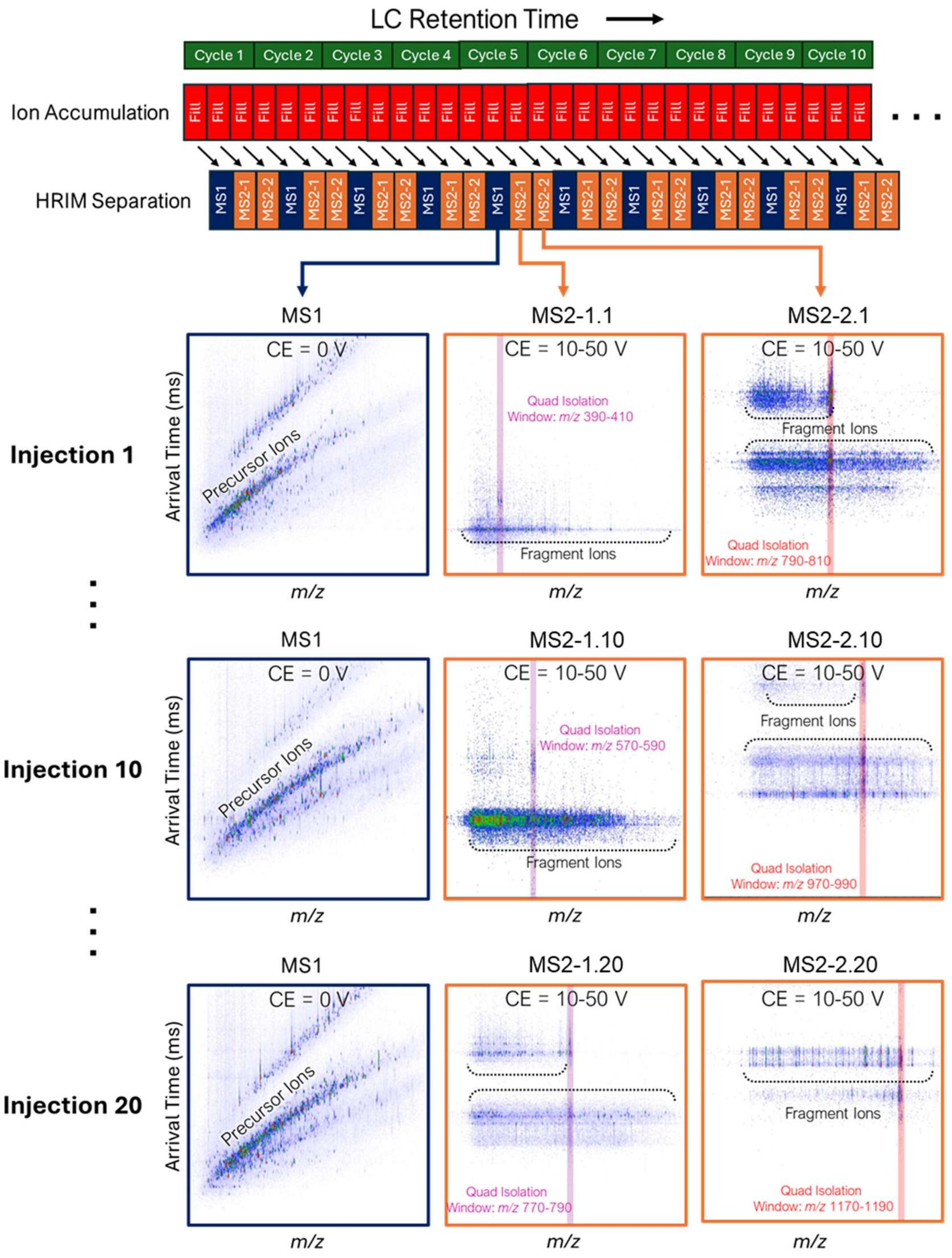


**Figure S3**: Schematic diagram of the pseudo-DIA-PAMAF workflow: A total of 20 injections were performed to cover the m/z range 390-1190 Th. Two MS scans were performed per cycle, with one quadruple isolation window per MS2 scan (i.e., MS2-1.1 and MS2-2.1 for injection 1, MS2-1.2 and MS2-2.2 for injection 2 and so on). The quadrupole isolation window width was set to 20 Th. Additional details are provided in Table S2

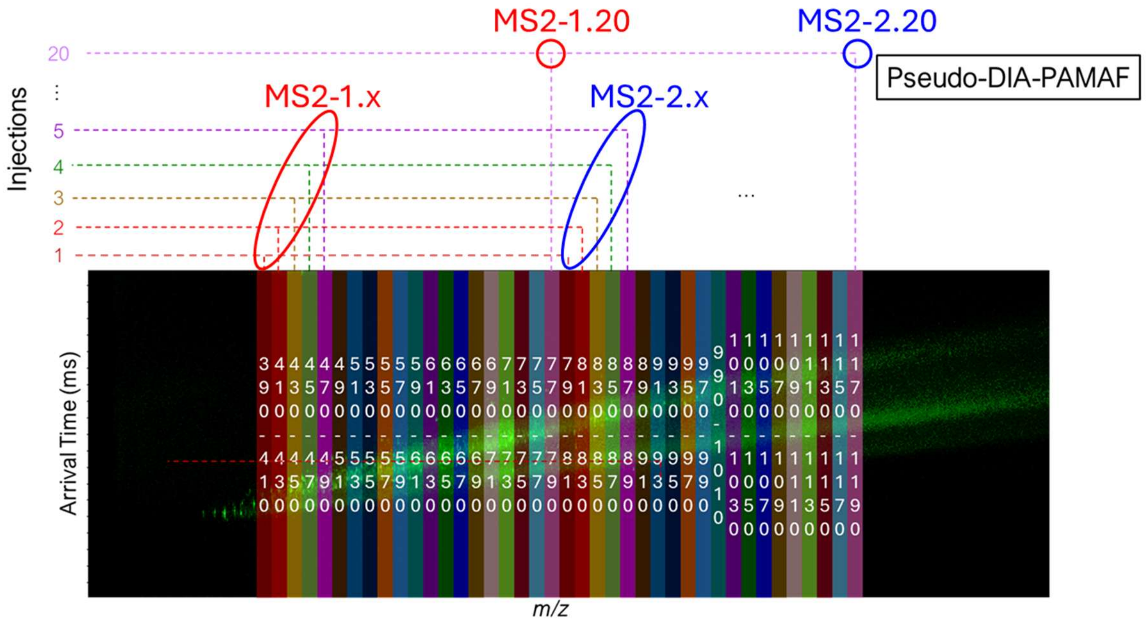


**Figure S4**: Overview of the quadrupole isolation windows used for pseudo-DIA-PAMAF mode experiments. Isolation windows are overlaid on the mobility vs. *m/z* heatmap.

**Table S2**: Parameters used for pseudo-DIA-PAMAF mode experiments.

| Parameter | Settings |
|---|---|
| *m/z* Range | 390-1190 Th |
| MS2 Acquisition | 2 MS2 scans per cycle |
| Isolation Strategy | One quadrupole isolation window per MS2 scan |
| Total Quadruple Windows | 40 |
| Quadrupole Isolation Window Width | 20 Th |
| Cycle Time | 1.325 s |
| Sample Load | 500 ng HeLa digest |
| OBA Fill Time | 390 ms |
| HRIM Frame Length | 405 ms |
| Number of Injections | 20 |

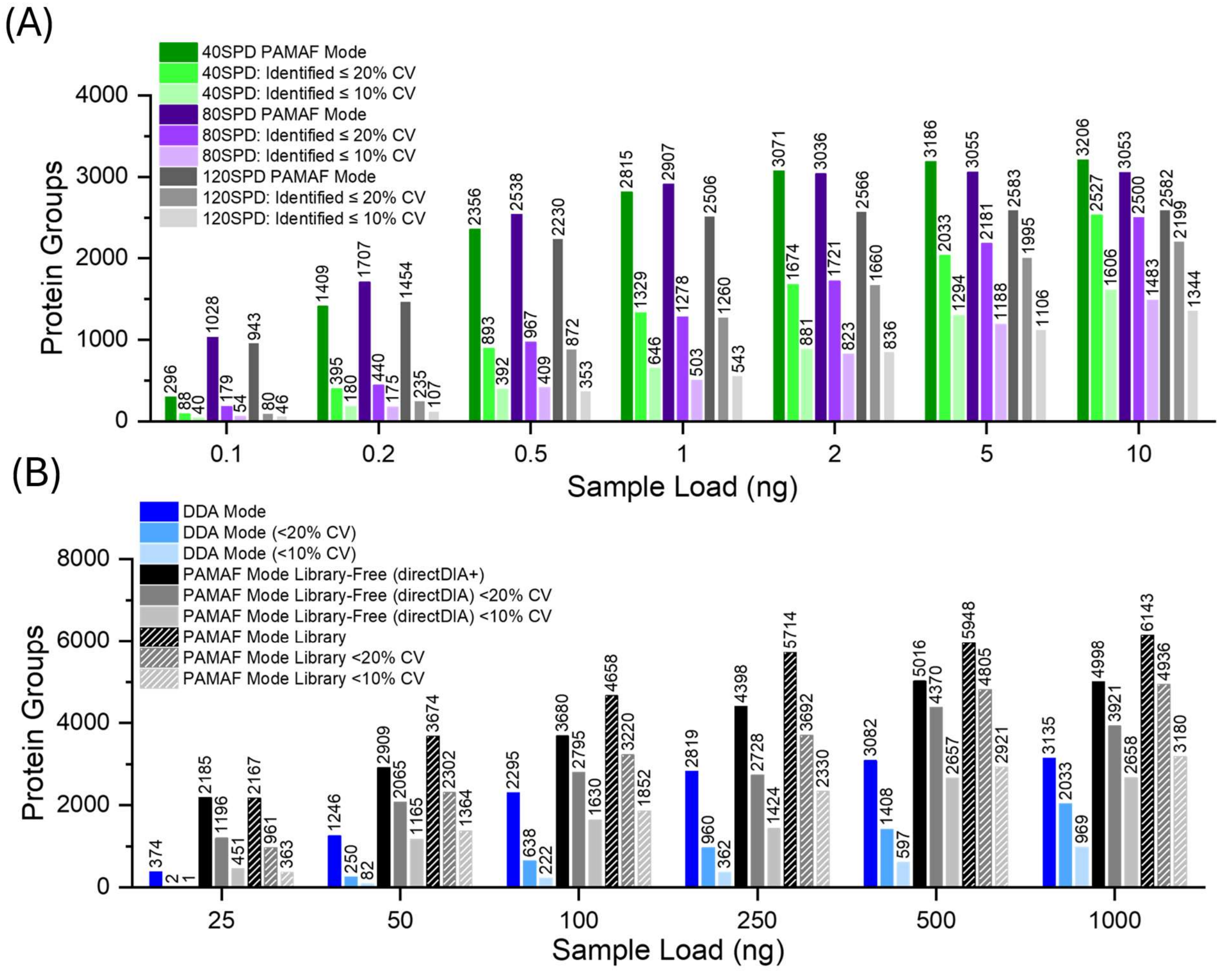


**Figure S5**: (**A**) Quantifiable protein groups from HeLa digests (low-load proteomics workflows, 100 pg - 10 ng) acquired in DDA and PAMAF modes at sample speeds of 40 SPD, 80 SPD, and 120 SPD. Data was processed in Spectronaut 19 using library-based search. (**B**) Quantifiable protein groups obtained from HeLa digests (high-load workflows, 25 – 1000 ng) using DDA, PAMAF, and pseudo-DIA-PAMAF modes at the same sample speed of 30 SPD. Data was processed in Spectronaut 19 using two distinct search strategies: library-free (directDIA+) and library-based search.
